# Computational Drug Design of Novel Agonists of the μ-Opioid Receptor to Inhibit Pain Signaling

**DOI:** 10.1101/2023.08.25.554876

**Authors:** Nancy Daoud, Diego Lopez Mateos, Mary A. Riley, Justin B. Siegel

**Affiliations:** Department of Chemistry, University of California Davis, Davis, California, United States of America; Department of Physiology and Membrane Biology, University of California Davis, Davis, California, United States of America; Biophysics Graduate Group, University of California Davis, Davis, California, United States of America; Microbiology Graduate Group, University of California Davis, Davis, California, United States of America; Genome Center, University of California Davis, Davis, California, United States of America; Department of Biochemistry and Molecular Medicine, University of California Davis, Davis, California, United States of America

## Abstract

Opioids such as Morphine, Codeine, Hydrocodone, and Oxycodone target the μ-opioid receptor, a G-protein-coupled receptor (GPCR), blocking the transmission of nociceptive signals. In this study, four opioids were analyzed for ADMET properties and molecular interactions with a GPCR crystal structure (PDB ID: 8EF6). This aided in the computational design of two novel drug candidates with improved docking scores and ADMET properties when compared to Hydrocodone. Homology analysis indicated that a *Mus musculus* (house mouse) animal model could be used in the preclinical studies of these drug candidates in the development of safer and more effective opioid drugs for pain management with reduced side effects.

## INTRODUCTION

Chronic pain is a serious health condition that leads to severe complications both physically and mentally.^1^ More than one in five adults experience chronic pain due to illnesses like cancer, fibromyalgia, and neuropathic pain, leading to more than 50 million adults experiencing long-lasting pain. Opioids are most often used in combination with anesthetics during surgical procedures to relieve pain and enhance sedation. They are also commonly prescribed to treat pain from serious injuries and post-surgery pain. Moreover, opioids play a crucial role in providing comfort and pain relief for patients in palliative care or at the end stages of life.^2^ Morphine, Codeine, Oxycodone, and Hydrocodone are four common opioids prescribed to treat such pain. They function by binding to opioid receptors on the nerve cells in the brain, spinal cord, as well as other parts of the body, and mimicking the natural painkiller endorphins produced by the brain.^3,4^ This results in opioids blocking pain signals to relieve moderate to severe chronic pain.

Opioids act on four types of G protein-coupled receptors (GPCRs) in the body, specifically, μ, δ, κ, and nociception receptors.^5,6^ These opioid receptors appear both on the cell bodies and axon terminals of neurons. Their protein structure consists of a long polypeptide chain that characteristically traverses the cell membrane seven times and couple with other inhibitory G-proteins intracellularly. GPCRs are activated by the binding of opioids extracellularly which alters the binding to the inhibitory intracellular G-proteins, thus mediating several cellular responses.

Prior to any opioid binding, the G-protein is inactive with a guanosine diphosphate (GDP) bound to the G-protein alpha (α), and beta gamma (βγ) subunits.^6^ When an opioid binds extracellularly, the GDP is released intracellularly, and guanosine triphosphate (GTP) binds in its place which releases the Gα subunit. The Gα subunit further separates from the Gβγ subunits, and both go on to directly interact with another G-protein gated potassium channel to keep it open. This lowers the electrostatic potential, decreases neuron excitability, blocks calcium from entering the cell, and ultimately prevents neurotransmitter release.^6^ These actions collectively contribute to the prevention of pain signaling.

However, even common opioid drugs such as Morphine, Codeine, Oxycodone, and Hydrocodone can have many negative side effects such as nausea, vomiting, constipation, dizziness, addiction, and mental fog.^2^ In this study, we aimed to improve these side effects by computationally designing two novel drug candidates with improved molecular interactions, docking scores, and ADMET properties than the current opioid drugs. Improving the binding affinity of these drugs would minimize side effects due to the decrease in off-target binding. The first novel drug candidate was designed using a drug discovery program while the second was by chemical intuition. A homolog analysis was also done to determine an animal model that could be used for preclinical studies based on the conservation of molecular interactions between the human and animal model. Overall, these novel candidates show great potential to be strong leads for improved drugs with less side effects over the current opioids on the market.

## METHODS

The crystal structure of Morphine-bound μ-opioid receptor-Gi complex (PDB ID: 8EF6) was obtained from RCSB Protein Data Bank website.^7^ The crystal structure was visualized in PyMOL to view ligand interactions and measure distances in Angstroms (Å).^8^

GaussView^9^ was used to build all drug compound structures, Gaussian09^10^ was used to optimize them, and OpenEye OMEGA^11^ was used to generate their conformer libraries. The visualization program VIDA^12^ was also used to analyze the drug compounds mentioned. OpenEye FILTER^13^ screened the molecules and determined the absorption, distribution, metabolism, excretion, and toxicity (ADMET) properties. ADMET reveals the pharmacokinetics and metabolic behaviors of these compounds based on their physical properties. This includes a Lipinski’s rule of 5 test which dictates that a drug compound must possess a molecular weight of less than 500g/mol, exhibit a XLogP value of less than 5, and have less than 5 hydrogen bond donors and 10 hydrogen bond acceptors. It also includes a filter test which determines if a drug has any toxicophores or toxic/reactive groups, and an aggregator test.

Make Receptor^14^ defined the binding site in the PDB crystal structure to be used for molecular docking. vBrood^15^ software was used with Hydrocodone to replace functional groups and generate bioisosteric compounds for potential new candidates. All drugs were docked into the active site using OpenEye and Fred^16^.

BLAST^17^ was employed to search for proteins homologous to the human Morphine-bound μ-opioid receptor-Gi complex (PDB ID: 8EF6) in animal models for preclinical studies. Then Jalview^18^ was used for the sequence alignment and to identify residues in these homologs that may interact with the novel drug candidates.

## RESULTS AND DISCUSSION

### Evaluation of four known opioid drugs

Four opioids (Morphine, Codeine, Hydrocodone, and Oxycodone) were analyzed and tested for ADMET properties and docking scores in the crystal structure of a human μ-opioid receptor-Gi complex (PDB ID: 8EF6). This control experiment aided in determining which opioid drug should be chosen for further optimization and creation into our two novel drug candidates.

The first evaluation for these four opioids was the ADMET screening (Table 1, Figure 1) where all four opioids fortunately passed the aggregator test and the Lipinski’s rule of 5 test. This suggests they all have a higher probability of exhibiting favorable ADMET properties. For example, they all exhibited a low XLogP value which suggests they have favorable oral bioavailability. Unfortunately for the filter test, only Hydrocodone and Oxycodone passed. This means that only Hydrocodone and Oxycodone do not possess any major toxicophoric groups.

**Table 1.** ADMET properties and docking scores of four opioid drugs.

| Name | Molecular Weight (g/mol) | Lipinski H-bond Donor | Lipinski H-bond Acceptor | XLogP | Number of Chiral Centers | Docking Score |
| --- | --- | --- | --- | --- | --- | --- |
| Morphine | 285.3 | 2 | 4 | 0.49 | 5 | -10.51 |
| Codeine | 300.4 | 1 | 4 | 1.23 | 5 | -9.62 |
| Hydrocodone | 299.4 | 0 | 4 | 1.51 | 4 | -9.40 |
| Oxycodone | 315.4 | 1 | 5 | 0.29 | 4 | -8.83 |

**Figure 1.**
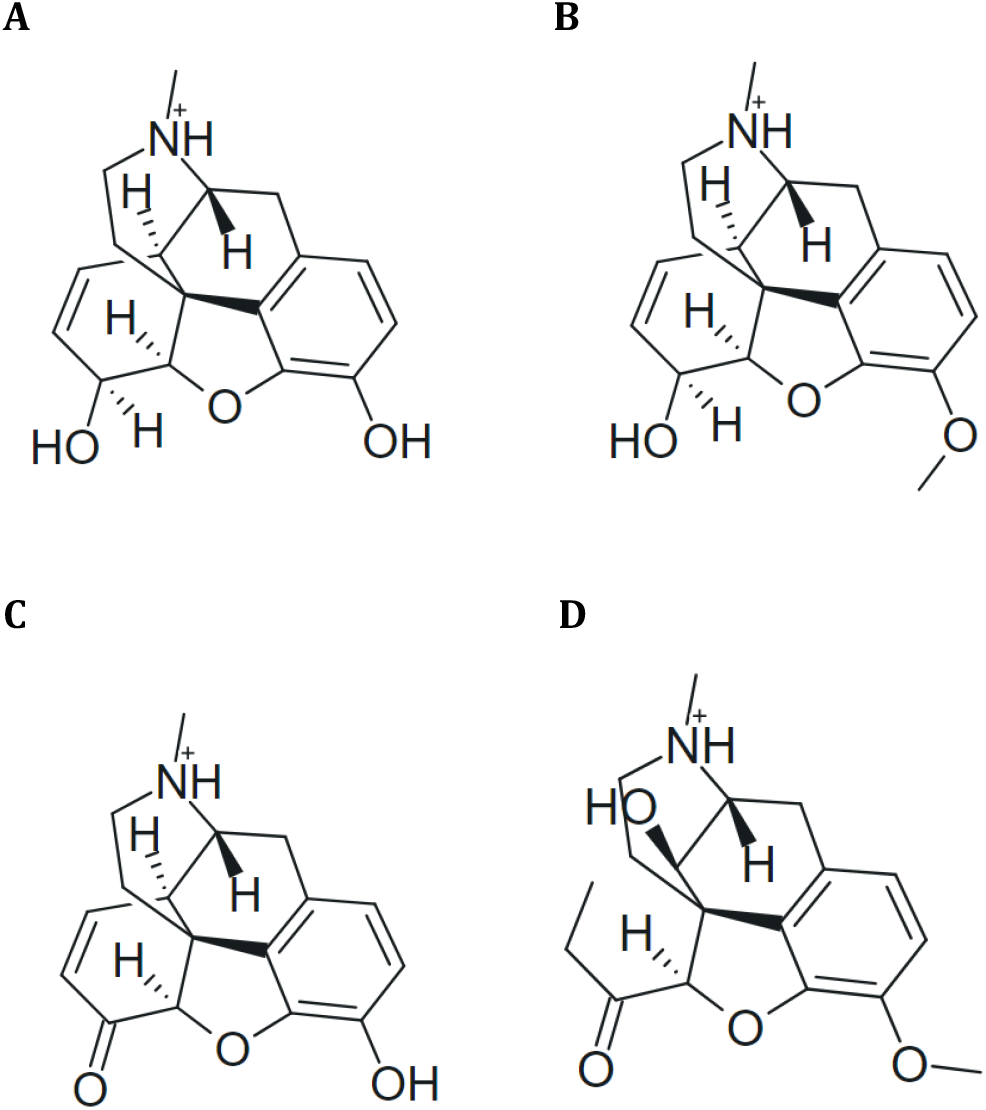
**A)** 2D structure of Morphine. **B)** 2D structure of Codeine. **C)** 2D structure of Hydrocodone. **D)** 2D structure of Oxycodone.

All four drugs have many chiral centers, each containing either four or five. Additionally, they all contain about the same number of hydrogen bond acceptors at around four or five. However, they differ in the number of hydrogen bond donors, with Morphine possessing the most at two, Codeine and Oxycodone with one, and Hydrocodone with none. This suggests Morphine may possess the most residue interactions after docking, while Hydrocodone may possess the least.

Next, each opioid was docked into the human μ-opioid receptor-Gi complex active site (PDB ID: 8EF6). The docking scores (Table 1) from highest to lowest were -10.50 for Morphine, -9.62 for Codeine, -9.40 for Hydrocodone, and -8.83 for Oxycodone. These scores suggest that there are no significant changes in binding energetics and thus, it is predicted to have similar binding affinity between all four opioids. Moreover, Morphine and Hydrocodone have the highest binding affinities for the active site, indicative of their more negative scores.

Overall, our findings in both ADMET scores and docking interactions suggest Morphine and Hydrocodone could both be good drugs to use to begin our designs. With both drugs containing the two highest docking scores and Hydrocodone having an ideal XLogP value of 1.51. Therefore, we chose to proceed with the opioid with one of the best docking score, the best XLogP value, and one of the few drugs to pass the filter test, Hydrocodone (Figure 1C).

First, the docked Hydrocodone opioid drug was assessed in PyMOL to determine its’ ligand-protein interactions (Figure 2A and 2B). PyMOL showed that Hydrocodone had a hydrogen bonding interaction with residue ASP 149 at a distance of 3.7Å, an electrostatic interaction with TYR 150 at 4.2 Å, and a pi-pi stacking interaction with TRP 295 at 4.0Å.

**Figure 2.**
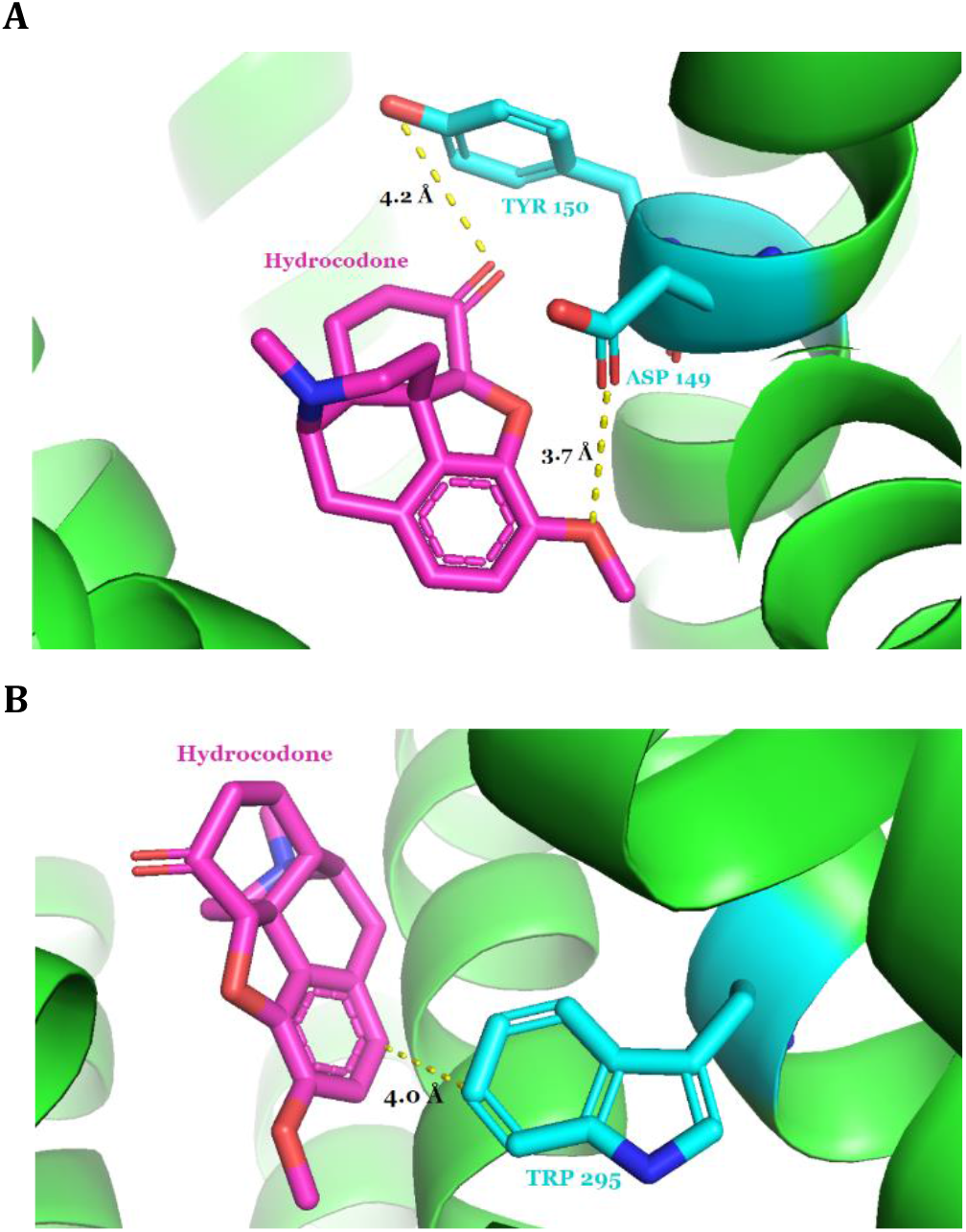
**A)** Hydrocodone (magenta) interactions with TYR 150 and ASP 149 residues (cyan). **B)** Hydrocodone (magenta) interactions with TRP 295 residue (cyan).

### Computationally Driven Design of Candidate 1

Candidate 1 (Figure 3A) was generated by loading Hydrocodone into vBrood for bioisosteric replacement modification of a methyl group to improve binding affinity. It possesses a newly protonated nitrogen atom as well as a cyclopropane attached to the oxygen atom to increase hydrophobicity. The generated molecule was analyzed using the same methods as for Hydrocodone and displayed a better docking score and ADMET properties than the original drug. This novel Candidate 1 drug has not been developed and was not found in SciFinder.

**Figure 3.**
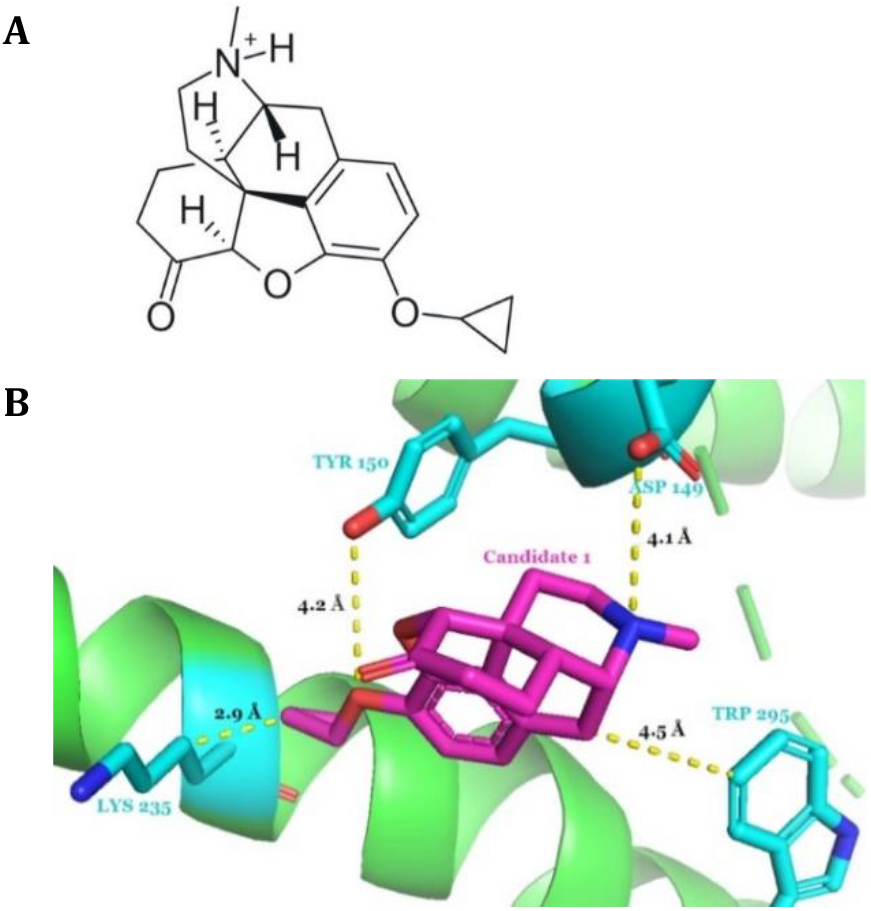
**A)** 2D structure of Candidate 1. **B)** Candidate 1 (magenta) interactions with ASP 149, TYR 150, TRP 295, and LYS 235 residues (cyan).

OpenEye FILTER was used to generate the ADMET properties for Candidate 1, shown in Table 2. Candidate 1 possesses a docking score of -13.49 which is a ∼39% improvement over Hydrocodone. The hydrogen bond score went from -0.15 to -4.40 in Candidate 1 (Table 3). This may be due to the increase in hydrogen bonding possibilities. It was also observed that protein and ligand desolvation both decreased from 2.41 to 2.00 and from 2.97 to 1.86, respectively, which are both good improvements. However, molecular weight increased and so did XLogP, but it’s still in range for good oral bioavailability. It also showed no violations in Lipinski’s Rule of 5.

**Table 2.** ADMET properties and docking scores of Candidates 1 & 2.

| Name | Molecular Weight (g/mol) | Lipinski H-bond Donor | Lipinski H-bond Acceptor | XLogP | Number of Chiral Centers | Docking Score |
| --- | --- | --- | --- | --- | --- | --- |
| Candidate 1 | 326.4 | 1 | 3 | 2.1 | 5 | -13.49 |
| Candidate 2 | 356.5 | 4 | 3 | 1.1 | 5 | -15.78 |

**Table 3.** Docking Analysis Scores of Hydrocodone, Candidate 1, and Candidate 2.

| Name | Shape Score | Hydrogen Bond Score | Protein Desolvation Score | Ligand Desolvation Score |
| --- | --- | --- | --- | --- |
| Hydrocodone | -14.63 | -0.15 | 2.41 | 2.97 |
| Candidate 1 | -12.98 | -4.40 | 2.00 | 1.86 |
| Candidate 2 | -14.66 | -6.29 | 2.41 | 2.76 |

The docking of Candidate 1 was then analyzed in PyMOL where it was noted that it could now reach more residues within the active site than the original drug. For example, Candidate 1 displays a new hydrophobic interaction between the cyclopropane and residue LYS 235 at 2.9Å (Figure 3B). This new interaction may be a contributor to the increased docking score. Candidate 1 partially retained an interaction with residue TRP 295 at a new distance of 4.5Å. However, the interaction involves a different ring in the structure and displays an increase from the 4.0Å distance with Hydrocodone. This most likely accumulates into a weaker interaction which was probably brought on by the altered docking pose when compared to the original drug.

Interestingly, the exact same electrostatic interaction between oxygen and residue TYR 150 in Hydrocodone exists at the same distance of 4.2Å in Candidate 1. This conservation suggests this may be an important interaction for binding. On the other hand, Candidate 1 shows an electrostatic interaction with residue ASP 149 at 4.1Å with a nitrogen atom that originally was a hydrogen bond with the original drug. This alteration may be due to the stereochemistry and pose conformation adjustments that occur with the structural changes. The general conservation of interactions with TRP 295, TYR 150, and ASP 149 residues suggests these may be important for any type of ligand binding. In review, even though Candidate 1 shows some weaker interactions, it forms new and altered interactions that overall increase its binding affinity over Hydrocodone.

### Chemical Intuition Driven Design of Candidate 2

Candidate 2 (Figure 4A) is a hypothesis-driven design based off Candidate 1, with an aim for an opioid with an improved docking score and ADMET properties. To do so, increased hydrogen bonding and reduced protein and ligand desolvation are ideal. Efforts resulted in replacing a few oxygen atoms with carbon atoms for increased stability. The double-bonded oxygen was replaced with a hydroxyl group and more hydroxyl groups were added for increased hydrogen bonding. Candidate 2 has not been developed and not found in SciFinder.

**Figure 4.**
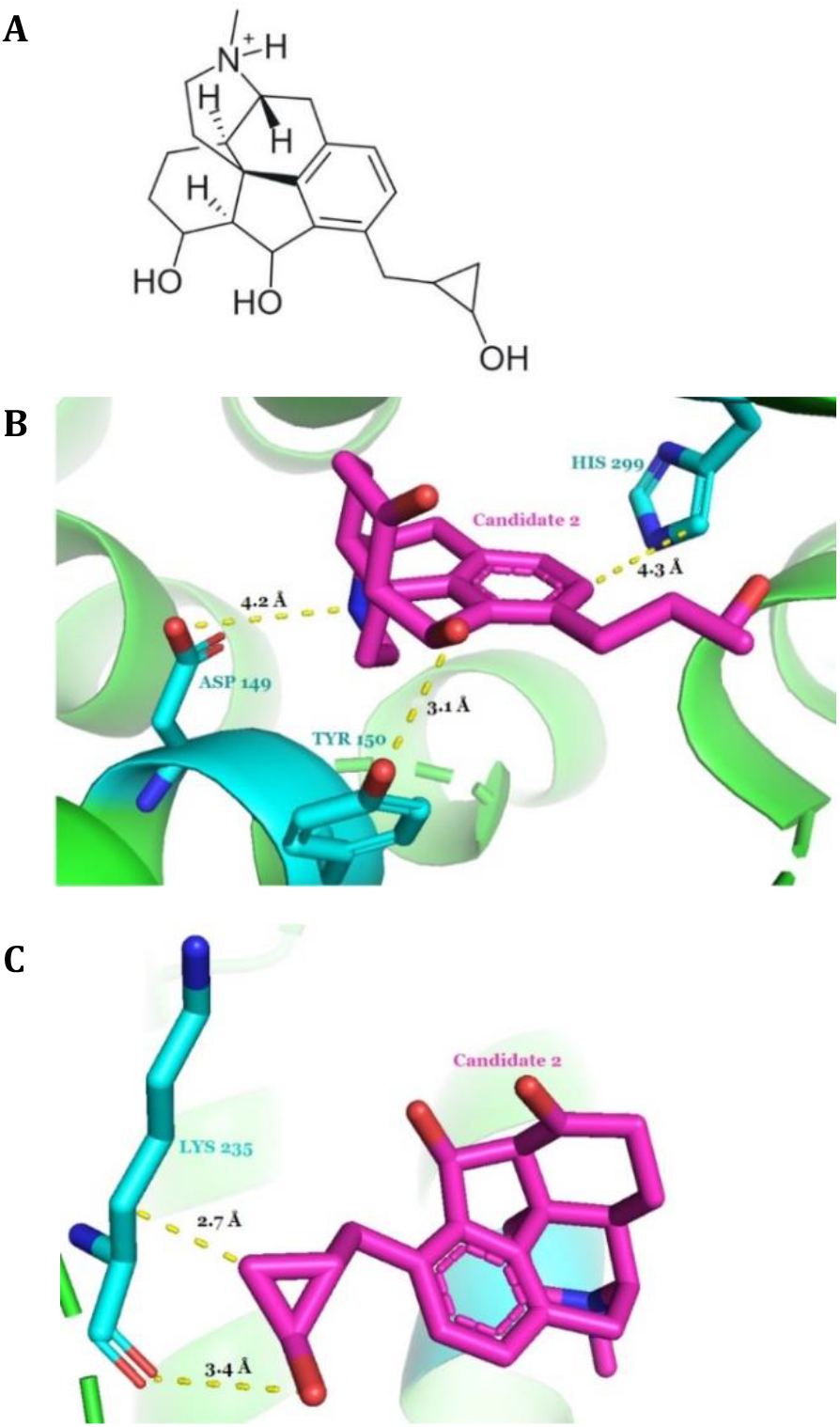
**A)** 2D structure of Candidate 2. **B)** Candidate 2 (magenta) interactions with ASP 149, TYR 150, and HIS 299 residues (cyan). **C)** Candidate 2 (magenta) interactions with LYS 235 residue (cyan).

The Candidate 2 design was then computationally built, optimized, docked into the opioid receptor, and underwent an ADMET screening. Results are shown in Tables 2 and 3. Analysis of Candidate 2 showed adding hydroxyl groups decreased the hydrogen bonding score from -4.40 in Candidate 1 to -6.29 in Candidate 2. More importantly, Candidate 2 had an even better docking score than Candidate 1, with a ∼63% improvement on Hydrocodone and a ∼17% improvement on Candidate 1. The XLogP is also a lot lower at 1.1, which suggests this drug will be absorbed, transported, and distributed in the body even better than Candidate 1. Overall, Candidate 2 shows both a better binding affinity and biochemical properties than Candidate 1 and the original Hydrocodone drug already on the market.

Candidate 2 was loaded into PyMOL to visualize any new interactions made (Figures 4B and 4C). An electrostatic interaction is seen between residue ASP 149 and Candidate 2 at a 4.2Å distance, which also occurs at a similar distance for Candidate 1 but was a stronger hydrogen bond interaction in Hydrocodone. Candidate 2 also retains an interaction with residue TYR 150 but is now altered to a stronger hydrogen bond at a distance of 3.1Å. That same interaction was previously only electrostatic and weak at 4.2Å in both Hydrocodone and Candidate 1.

Interestingly, Candidate 2 shows a new pi-pi stacking interaction with residue HIS 299 at a 4.3Å distance with which no previous drugs seem to have any interaction. The cyclopropane shows the same hydrophobic interaction with residue LYS 235 that occurs in Candidate 1, but at a stronger shorter distance of 2.7Å. Moreover, a new hydrogen bond is made with LYS 235 at 3.4Å (Figure 4C). This is a new interaction resulting from the added hydroxyl group on the cyclopropane.

Overall, Candidate 2 which was a chemical-intuition driven drug design shows much promise with its improved docking score, hydrogen bonding interactions, and lower XLogP number when compared to the original Hydrocodone drug.

### Homology Analysis for Preclinical Animal Testing

Clinical trials are an important step to examine the viability of novel drug candidates to become potential drugs on the market for chronic pain. Based on a BLAST search, it was discovered that a few organisms contain homologous proteins to the human μ-opioid receptor protein sequence. *Mus musculus* (mouse) have a 100% query cover and a high percentage identity of 97.80% homolgous protein. *Mus musculus* is a very common starting organism to test due to their physiological and genetic similarity to humans.

Jalview was used next to do an in-depth protein sequence alignment analysis to evaluate if the drug candidates would interact with the homologous protein active site in the same way as it would in humans. The sequence alignment showed that ASP 149, TYR 150, TRP 295, and LYS 235 residues that interact with both Candidate 1 and Candidate 2 were all highly conserved. Additionally, the residue HIS 299 which only interacts with Candidate 2 was also highly conserved. This suggests the mouse homolog protein will likely interact and bind similarly to both Candidate 1 and Candidate 2 as it would in a human. Therefore, the *Mus musculus* animal model should work in further testing needed to determine the safety and side effects of these novel drugs.

Furthermore, beneficial microbes and pathogenic microbes were also searched for on BLAST to see if they have any homologs to the target protein causing off-target effects. After several tries with multiple classes and species, there were no matches. This suggests that Candidates 1 and 2 should not have any effect on them.

## CONCLUSION

Chronic pain is a common and serious health condition that affects millions of people every year.^1^ The opioids existing in the market need improvement due to the many side effects and other risks such as overdose and dependence.^2^ Developing new opioid medicine that has better absorption, distribution, metabolism, and less toxicity is crucial. As well as drugs with fewer off-target interactions. We aimed to do this by modifying an already known drug, Hydrocodone, with computational and chemical intuition methods. We thus proposed two novel drug candidates with great potential showing improved ADMET and docking scores over the original drug. These candidates still need to be synthesized and tested, ideally on a *Mus musculus* animal model, to find how they behave in a biological setting and determine their safety for the public.

## AUTHOR INFORMATION

### Author Contributions

Research was designed by all authors; all experiments were carried out by N.D. The manuscript was written through contributions of all authors. All authors have given approval to the final version of the manuscript.

## ACKNOWLEDGMENT

Research reported in this publication was supported by UC Davis, the National Science Foundation Award Numbers 1827246, 1805510, 1627539, the National Institute of Environmental Health Sciences of the National Institutes of Health (NIH) under Award Number P42ES004699, UC Davis, NIH Award Number R01 GM 076324-11, and the Rosetta Commons. The content is solely the responsibility of the authors and does not necessarily represent the views of the National Institutes of Health or National Science Foundation. This study was derived from a course based undergraduate research study conducted in Chemistry 130B at UC Davis.

